# Antibody and T cell memory immune response after two doses of the BNT162b2 mRNA vaccine in older adults with and without prior SARS-CoV-2 infection

**DOI:** 10.1101/2021.07.08.451426

**Authors:** Julie Demaret, Bénédicte Corroyer-Simovic, Enagnon Kazali Alidjinou, Anne Goffard, Jacques Trauet, Sophie Miczek, Fanny Vuotto, Arnaud Dendooven, Dominique Huvent-Grelle, Juliette Podvin, Daniel Dreuil, Karine Faure, Dominique Deplanque, Laurence Bocket, Alain Duhamel, Julien Labreuche, Annie Sobaszek, Francois Puisieux, Myriam Labalette, Guillaume Lefèvre

## Abstract

We quantified S1-specific IgG, neutralizing antibody titers, specific IFNγ secreting T cells and functionality of specific CD4+ and CD8+ T cells in 130 young adults (median age 44.0 years) and 106 older residents living in a long-term care facility (86.5 years) after 2 doses of BNT162b2. Three months after the first injection, humoral and cellular memory responses were dramatically impaired in the 54 COVID-19-naive older compared to the 121 COVID-19-naive younger adults. Notably, older participants’ neutralizing antibodies, detected in 76.5% (versus 100% in young adults, *P* < 0.0001), were ten times lower than the younger’s antibody titers (*P* < 0.0001). Antibody and T cell responses were greater among the 52 COVID-19-recovered than among the 54 COVID-19-naive older adults (*P* < 0.0001). Our study shows that 2 doses of BNT162b2 does not guarantee long-term protection against SARS-CoV-2 in the older. An additional dose should be considered to boost their specific memory response.

## Main

Since the emergence of severe acute respiratory syndrome coronavirus 2 (SARS-CoV-2) and the beginning of the worldwide coronavirus disease 2019 (COVID-19) pandemic, unprecedented efforts have been made to develop vaccines. Considered among the most at risk of developing severe COVID-19, long-term care facility (LTCF) older residents were among the first to be vaccinated. In addition to age, older adults usually cumulate other risk factors for COVID-19 and death, including diabetes, hypertension, cardiovascular disease and/or malignancy (1). Furthermore, the closed environment and the relative inability of residents to adopt preventive health measures led to numerous outbreaks in LTCFs worldwide (1). For these reasons, there were high hopes for anti-SARS-CoV-2 vaccines, especially among the older and health-care workers (HCW) in LTCFs. However, the older display physiological alterations of cellular and humoral immunity that affect vaccine responses (2,3), and, due to their age and frailty, they were not included in clinical trials evaluating the BNT162b2 mRNA vaccine (4–7).

The aim of this study was to assess the specific memory humoral and cellular response to the BNT162b2 mRNA vaccine in older LTCF residents. At the time of this study, French health authorities still recommended the two-dose vaccination regimen for LTCF residents who presented with COVID-19 in the months prior. Consequently, in order to evaluate the impact of prevaccine immunization, we compared the post-vaccinal response in COVID-19-naive and COVID-19-recovered older LCTF residents (hereafter referred to as older residents or older adults).

For all analyses, we focused on the S1 domain of the SARS-CoV-2 spike protein, which bears the receptor binding domain (RBD). RBD is of major interest since it enables SARS-CoV-2 to bind to the ACE2 receptor on targeted cells. Specific humoral response was assessed by serum anti-S1 IgG titers and by a neutralization assay based on live SARS-CoV-2 (LV-NT) and a pseudovirus-based neutralization (pV-NT) assay. Considering the major role of the T cell response in older adults, we quantified specific T cells using S1 peptide pools to stimulate specific cells which were detected by interferon-gamma (IFNγ) release assay (ELISpot) (8) and surface activation-induced markers (AIM). The functionality of specific CD4+ and CD8+ T cells was assessed by intracellular IFNγ, interleukine-2 (IL-2) and tumor-necrosis-factor alpha (TNFα) production by flow cytometry (Extended Data Fig. 1).

We consecutively included 130 HCWs (hereafter referred to as young adults) (median [interquartile range, IQR] age 44.0 [39.7;50.5] years) and 106 older residents (median [IQR] age 86.5 [81.0;90.0] years) who had received two vaccine doses between December 2020 and April 2021, with no known comorbidities and who did not receive any concomitant treatment that could attenuate the immune response (Extended Data Fig. 2 and Supplementary Table 1). All participants were sampled before (D0) and 90 days (D90) after the first dose.

Our primary objective was to compare the specific memory response in COVID-19-naive older (*n =* 54) and in COVID-19-naive younger adults (*n =* 121). At D0, no participant in either group had detectable anti-S1 IgG antibodies or neutralizing antibodies. Some participants had exceptionally low counts of S1 peptide pools reactive T cells detected by ELISpot (median [IQR] 1 [0;2] cell/250.000 peripheral blood mononuclear cells in both groups) (Extended Data Fig. 3a,b). At D90, S1 IgG reactivity was detected in almost all participants in both groups (99.2% of younger and 97.2% of older adults), but the median titer of anti-S1 IgG antibodies was 2 times lower among the older residents (*P* < 0.001) (Fig. 1a). The difference was greater for neutralizing antibodies, with a geometric mean of 50% serum neutralization titer (NT50) 10.2 times lower in the older group according to the LV-NT assay (Fig. 1b) (mean [95% confidence interval, CI] 29.8 [16.0;55.2] versus 305.0 [243.1;382.6]; median [IQR] titers: 40.0 [5.7;160.0] versus 320.0 [160.0;640.0], *P* < 0.0001). The number of responders (*ie,* participants who had detectable neutralizing antibodies) was 39 (*n* = 51 available data, 76.5%) COVID-19-naive older adults and 101 (*n =* 101 available data, 100%,) (*P* < 0.0001) COVID-19-naive younger adults (Fig. 1c). The mean NT50 in each group was consistent with the pV-NT50 assay (Fig. 1d).

**Fig. 1:**
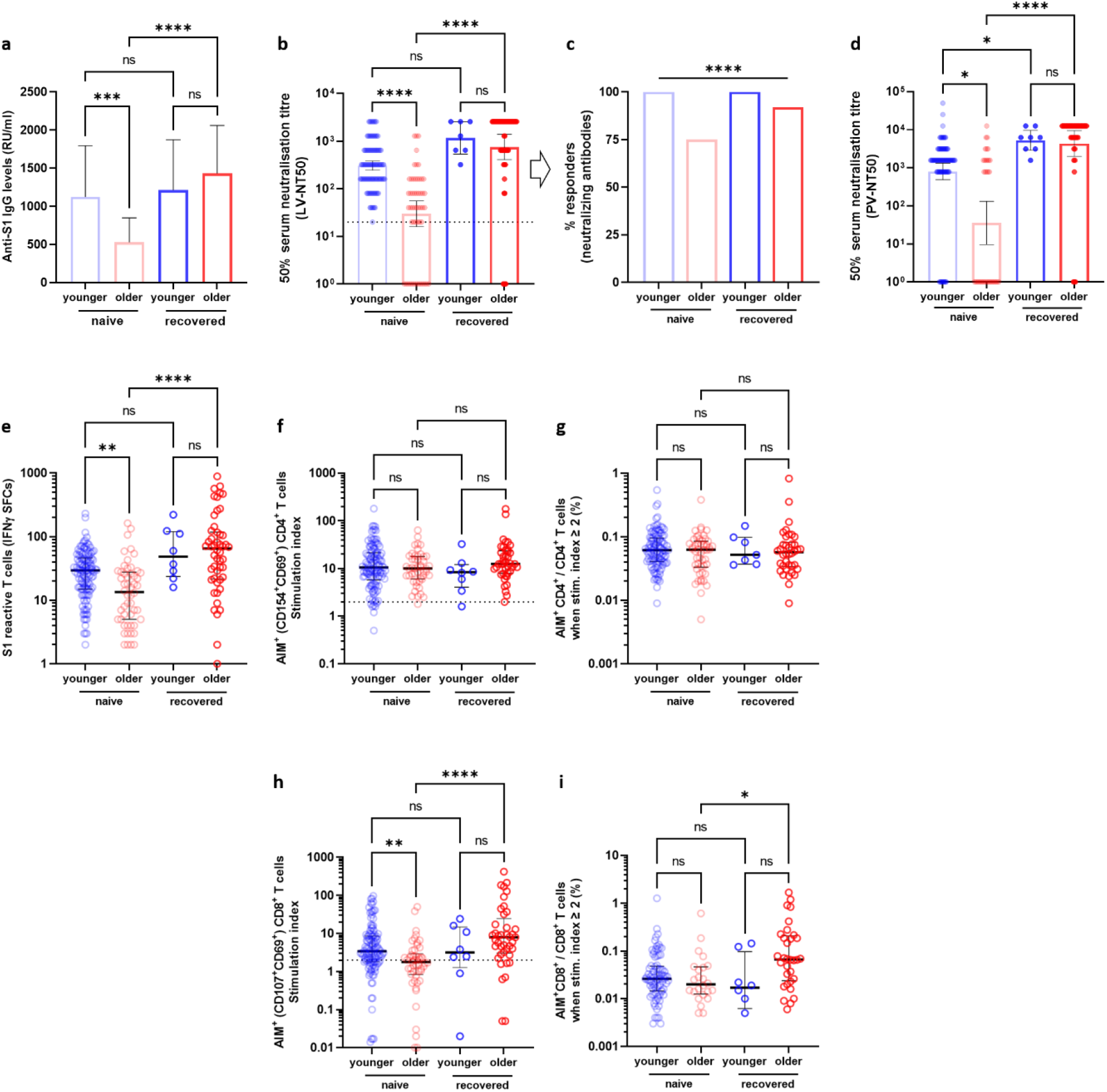
Specific antibody and T cell responses in older and in young adults 3 months after the first injection of BNT162b2. **a-d.** Antibody responses assessed by ELISA (anti-S1 IgG) (COVID-19-naive young adults *n* = 121, COVID-19-naive older *n* = 54, COVID-19-recovered young *n* = 8, COVID-19-recovered older *n* = 47; median [interquartile range, IQR] are shown); **a.** serum neutralization assay against live virus (COVID-19-naive young adults *n* = 101, COVID-19-naive older *n* = 52, COVID-19-recovered young *n* = 7, COVID-19-recovered older *n* = 51; geometric median and 95% confidence interval are shown); **b**. participants with detectable neutralizing antibodies according to live virus neutralizing assay (titer ≥ 1:20); **c.** and serum neutralization assay against pseudovirus (COVID-19-naive young adults *n* = 103, COVID-19-naive older *n* = 36, convalescent control *n* = 8, convalescent elderly *n* = 41; geometric median and 95% confidence interval are shown); **d,e.** number of S1 peptide pool reactive T cells (ELISpot) (COVID-19-naive young adults n=121, COVID-19-naive older *n* = 52, COVID-19-recovered young *n* = 8, COVID-19-recovered older *n* = 50; median [interquartile range, IQR] are shown). **f-i.** Specific CD4^+^; **f,g.** and CD8^+^; **h,i.** T cells according to activation induced markers (AIM), reported with their stimulation index; **f-h**. percentage of responders, ie participants with a stimulation index ≥2; **g-i.** COVID-19-naive young adults *n* = 113, COVID-19-naive older *n* = 48, COVID-19-recovered young *n* = 8, COVID-19-recovered older *n* = 41; median [interquartile range, IQR] are shown. *P* values * < 0.05, ** < 0.01, *** < 0.001, **** < 0.0001, ns: not significant. AIM^+^, cell expressing activation induced markers; IFNγ SFCs, interferon gamma spot forming cells; LV-NT50, 50% serum neutralization titer in live virus neutralization assay; pV-NT50, 50% serum neutralization titer in pseudovirus neutralization assay. Kruskal-Wallis test (with *post-hoc* Dunn) were used for multiple comparisons, Fisher’s exact test was applied for comparisons of responder frequency.

Regarding cellular response, T cells reactive to the S1 subunit detected by ELISpot were less frequent in the older than in the younger group (29.5 [15.0;46.5] versus 13.5 [25.0-27.57], respectively) (*P* = 0.002) (Fig. 1e). The distinction of S1-specific CD4+ and CD8+ T cells was studied by quantification of cells expressing the surface activation-induced markers (AIM+ T cells) (Extended Data Fig 4a). We observed no significant difference in the rate of participants with detectable AIM+ CD4+ T cells between the COVID-19-naive young and old adults (89.8% versus 97.9%, respectively, *P* > 0.05). The frequency of AIM+ CD4+ among CD4+ T cell counts were also similar in both groups (Fig. 1f,g). Conversely, more COVID-19-naive young participants developed AIM+ CD8+ T cells than COVID-19-naive older adults (76.4% versus 48.0%, respectively, *P =* 0.0018), but the frequency of AIM+ CD8+ among total CD8+ T cells was similar in both groups (Fig. 1h,i).

Finally, we analyzed all acquired parameters in COVID-19-naive young and older participants. Age negatively correlated with anti-S1 IgG, neutralizing titers, count of specific IFNg secreting T cells (ELISpot), and CD4 and CD8+ AIM+ T cells, which supports the differences observed between the two groups. There were also strong correlations between the immune parameters, which highlights both the conserved links between these different adaptive responses among the older population, and the robustness of the chosen approaches (Extended Data Fig. 5 and Supplementary Table 2).

Our secondary objective was to evaluate the capacity of the vaccination to boost the natural anti-SARS-CoV-2 memory response. We compared the 3-month post-vaccinal response in COVID-19-naive and in COVID-19-recovered older residents. Among the included participants, 51 COVID-19-recovered older adults (*n* = 5 according to high Anti-S1 IgG titers, *n* = 46 by positive PCR: median [IQR] interval 4.2 [3.3-8.3] months) were compared to COVID-19-naive counterparts. Only eight COVID-19-recovered young participants were available, which limited the possible comparisons among the naive and recovered young adults. We observed that older participants with prior COVID-19 had lower baseline anti-S1 IgG levels (*P* < 0.0001) and neutralizing antibodies (*P* = 0.04) than the COVID-19-naive counterparts after 2 doses of BNT162b2 mRNA vaccine, and similar S1 reactive T cells detected by ELISpot (Extended Data Fig. 6). Taken together, the data suggest that, among the older population, the vaccine-induced antibody response may be better than natural immunization, as previously observed in the phase 2 trial evaluating BNT162b immunogenicity (5). Furthermore, the anti-S1 IgG (Fig. 1a) and the neutralizing antibody levels (Fig. 1b,c,d), IFNγ secreting T cell counts (ELISpot) (Fig. 1e) and AIM+ CD8+ T cells (Fig. 1h,i) after 2 vaccine doses increase to a greater extent among COVID-19-recovered older adults compared to COVID-19-naive older adults. In addition, 92.2% of COVID-19-recovered older adults produced detectable neutralizing antibodies (Fig. 1c) compared to 76.5% of COVID-19-naive older participants. These results suggest that this vaccination is highly efficient in boosting the prior natural memory response. We believe this message is particularly compelling: while the progressive decrease of specific immune response after COVID-19 has been reported (15), many argue that COVID-19 exposure is provides sufficient immunity and that vaccination could be unnecessary or even detrimental for COVID-19-recovered older adults.

Our study also brings to light new elements on the function of T cells in the older population after 2 doses of BNT162b2. To better assess the T cell response in COVID-19-naive and COVID-19-recovered older adults, we studied the cytokine production. The percentage of cells able to produce 1, 2 and/or 3 cytokines among INFγ, IL-2 and TNFα (polyfunctional cells) differed according to age and past history of COVID-19 exposure (Fig. 2a). Indeed, the amount of specific IFNγ+ and IFNγ+IL-2+TNFα+ (triple+) CD4+T cells were lower in COVID-19-naive older participants than in COVID-19-naive young adults, but were over 3 times greater in the COVID-19-recovered than in the COVID-19-naive older participants (Fig. 2b). Conversely, TNFα+ cells were more frequent among specific CD8+ in COVID-19-recovered compared to COVID-19-naive older adults (Fig. 2c).. We also noted that, in the COVID-19-naive population, the frequency of AIM+IL-2+CD8+ T cells was greater among the older than among the younger. This suggests that, while CD8+ T cells can be highly activated with SARS-CoV-2 antigens in the naive older, the main effector cytokines required for antiviral response are not produced (Fig. 2c).

**Fig. 2:**
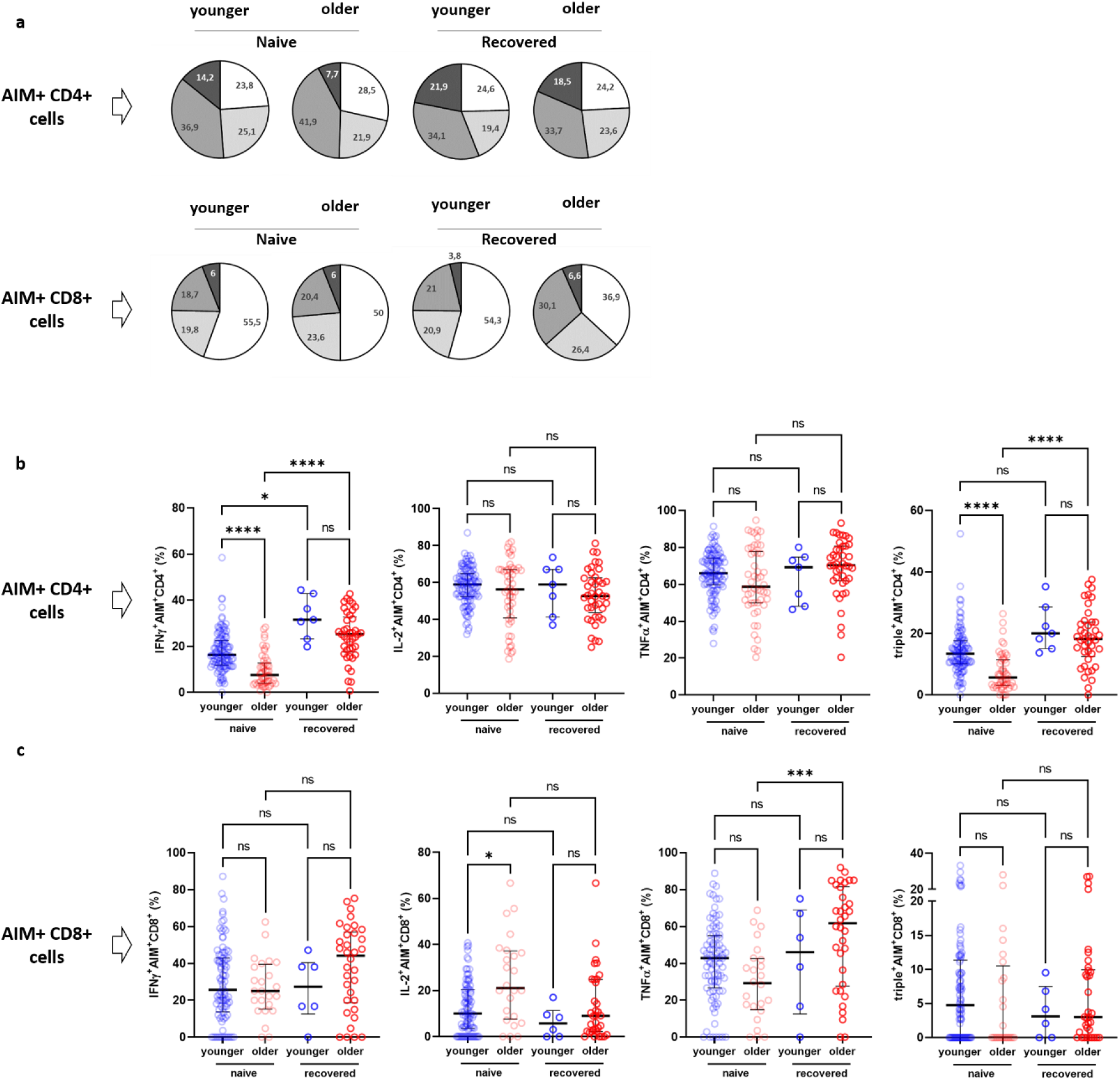
Functionality of specific CD4+ and CD8+ T cells in elderly LTCF residents and HCWs (control group) 3 months after the first injection of BNT162b2. **a.** Pie charts representing the relative proportions of AIM+ CD4+ and CD8+ T cells producing none (white), one (light grey), two (medium grey) or three cytokines (dark grey) out of INFγ, IL-2 and TNFα according to participant group and past history of COVID-19 (naive and recovered); **b.** Proportion of AIM+ CD4+ cells producing IFNγ, IL-2, TNFα and proportion of IFNγ+IL-2+TNFα+ (triple+) CD4+ T cells according to participant group and past history of COVID-19 (naive and recovered). (COVID-19-naive young adults *n* = 113, COVID-19-naive older *n* = 48, COVID-19-recovered young *n* = 8, COVID-19-recovered older *n* = 41; median [interquartile range, IQR] are shown) **c.** Proportion of AIM+ CD8+ cells producing IFNγ, IL-2, TNFα and proportion of IFNγ+IL-2+TNFα+ (triple+) CD8+ T cells according to participant group and past history of COVID-19 (naive and recovered) (COVID-19-naive young adults *n* = 113, COVID-19-naive older *n* = 48, COVID-19-recovered young *n* = 8, COVID-19-recovered older *n* = 41; median [interquartile range, IQR] are shown). *P* values * < 0.05, ** < 0.01, *** < 0.001, **** < 0.0001, ns, not significant. AIM+, cell expressing activation induced markers. The Kruskal-Wallis test (with *post-hoc* Dunn) was used for multiple comparisons.

To confirm the significance of our results in specific cellular responses, we used an automatized cluster analysis of T cell subsets (Extended Data Fig. 7). The hierarchical clustering confirmed the lower cytokine production in COVID-19-naive older participants compared to the three other groups of interest. The cluster analysis also showed a higher CD8+/CD4+ ratio among AIM+ T cells in the COVID-19-recovered older group. Similarly, an unsupervised analysis using t-distributed stochastic neighbor embedding (t-SNE) confirmed the lower cytokine production in COVID-19-naive compared to the COVID-19-recovered older participants (Extended Data Fig. 8). Indeed, specific IFNγ+ and polyfunctional CD4+ T cells, the most important ones in the orchestration of the whole adaptative immune response, decreased in older adults (compared to the younger participants). As these poor specific responses and specific TNFα+ CD8+ T cells, the most important in antiviral defense, were also highly boosted in participants who had a prior COVID-19 infection, a repeated vaccination could be effective in increasing immune effector cells in the older population. This is particularly important if we consider the effector roles of these cells in limiting the disease severity (16,17)

We evaluated whether some classical immunosenescence parameters could account for the poor vaccinal response among the older adults. As other authors, we failed to identify any link between vaccinal response and frailty (9), nutritional state (except between serum albumin levels and neutralizing antibodies with pV-NT assay) (Extended Data Fig. 9), or baseline T cell and naive T cell counts (Extended Data Fig. 10). A positive correlation was observed between baseline B cell count and neutralizing titers, although only with the LV-NT assay (Extended Data Fig. 10). We also assessed plasma levels of IL-1β, IL-6, TNFα and IL-10 in COVID-19-naive older adults. Plasma IL-1β levels tended to be negatively correlated with anti-S1 IgG and live virus neutralizing antibodies, and plasma TNFα levels correlated negatively with both neutralizing titers (Extended Data Fig. 11). These data suggest that “inflammageing” may play a role in the poor antiSARS-CoV-2 antibody response in the older. This is in line with previous data in human and mice models, which reported that serum TNFα negatively correlated with the B cell response and a vaccine-specific antibody response (10).

Overall, our work demonstrates that COVID-19-naive older adults have a poor memory immune response to BNT162b2 mRNA vaccine compared to the younger adults. Our results are in line with earlier evaluations of antibody and/or T cells response after a first dose (9,11) and between 14 to 22 days after the second one (9,12,13), indicating a poorer response in the elderly (10,11). Even though we did not assess memory switched B cells or memory phenotype T cells, our results obtained 90 days after the first dose and 60 days after the second dose may reflect the memory response established after vaccination rather than a response being initiated and may predict that immunity may wane even more over time.

Our study also demonstrated that specific memory response is greater in COVID-19-recovered older residents (compared to COVID-19-naive), and at a level similar to that of young participants. These results are in agreement with previous reports suggesting that patients with prior COVID-19 infection had a better antibody response, regardless of the age (9,13,14). Our study illustrates that, even if the ability to respond to neoantigens is usually impaired in the older, the post-COVID-19 memory immune response is improved by an additional boost. Our study is, however, limited in that, due to the recommendations applied in France at the time of the study, we were unable to assess whether a single dose of this vaccine after exposure to COVID-19 would have generated a sufficient response in these individuals. Also, the relatively short follow-up period only allowed us to assess the short-term effects of the vaccine. However, these data on the 2-month residual immune memory after the second dose may help anticipate future needs to adapt the vaccination strategy among the older.

To conclude, our results demonstrate that, with the recommended vaccination scheme (*ie,* 2 doses of BNT162b2), both antibody and cellular responses are impaired in the COVID-19 naive older population compared to the younger group. Our work also shows that the post COVID-19 memory response can be boosted by two doses of BNT162b2, and that a 3-dose instead of a 2-dose strategy may be relevant among the COVID-19 naive older population. Additionally, HCWs in LTCFs should be aware of the importance of being vaccinated themselves. Further work will include post-vaccinal monitoring to assess the outcome of post-vaccinal response at 9 months in both older and young participants.

## Acknowledgements

We thank Dominique Becuwe, Véronique Betrancourt, Virginie Dutriez, Anne Guigo, Coralie Lefebvre, Véronique Lekeux, Marie-Thérèse Meleszka and Catherine Mortka for their technical support and Bertrand Accart, Michael Hisbergues and Camille Tognelli for their contribution (Centre de Ressource Biologique). We also thank Séverine Duflos, Marie Broyez, Peggy Bouquet, Clémentine Roland, Marion Lecorche, Abeer Shaikh Al Arab, Isabelle Tonnerre, Japhete Elenga Koanga, Laurent Schwarb Emilie Rambaut, and all the nurses implicated in patients sampling, and Sarah Abdesselam and François Rose for data collection. Our thanks also to Melissa Charbit and Jone Iriondo (Excelya) for help in reviewing and editing the manuscript. This work was supported by the French government through the Programme Investissement d’Avenir (I-SITE ULNE/ANR-16-IDEX-0004 ULNE) managed by the Agence Nationale de la Recherche.

## Author contributions

JD, BCS, ML and GL conceived and designed the study and participated in data collection, analysis, writing of the manuscript and revision of the manuscript.

EKA, AG, JT, SM, FV, AD, DHG, JP, DD, KF, DD, LB, AD, JL, AS and FP participated in data collection, analysis and revision of the manuscript.

## Competing interests

None of the authors have any competing financial or non-financial interests.

## Data availability statement

The datasets generated during and/or analyzed during the current study are available from the corresponding author on reasonable request.

## SUPPLEMENTARY FILES

### METHODS

#### Study design and participants

This was a prospective single-center study conducted at the Lille University Hospital, in the North of France. Participants were consecutively included in the study, and were healthcare workers (HCW; hereafter referred to as young adults) aged 18-65 years and long-term care facility (LCTF) residents (hereafter referred to as older residents or older adults) aged >65 years who consented to be vaccinated with BNT162b2 mRNA vaccine and were willing to comply with the study procedures. None of the enrolled participants had a recent, current or persistent infectious disease, any neoplasia diagnosis in the last 5 years, or treatment with steroids and/or immunosuppressants. Participant characteristics collected at baseline included confirmation of prior SARS-CoV-2 infection, determined by polymerase chain reaction (PCR) and/or high antibody titer to SARS-CoV-2 spike S1 domain: participants with a history of positive PCR and/or who tested positive for anti-S1 antibodies were considered as COVID-19-recovered, and the other participants as “COVID-19-naive”. Among older adults, Geriatric Nutritional Risk Index was calculated according to the Lorentz formula: GNRI = (1.489 × albumin, g/l) + (41.7 × present/ideal body weight), with the ideal weight calculated according to the Lorentz formula (1). Frailty was assessed with the Clinical Frailty Scale as proposed by Rockwood et al. (2), and using the Fried frailty phenotype criteria (3). All participants received the two-dose BNT162b2 vaccination at a 3-week dosing interval: the first dose was administered at Day 0 (D0), and the second dose between D21 and D28. Serum samples were collected for all participants at D0, and D90 (±14 days) after the first dose.

#### Anti-SARS-CoV-2 antibodies

Anti-SARS-CoV2 spike S1 domain-specific Immunoglobulin G (IgG) was assessed in serum samples using ELISA (Quantivac, Euroimmun Lübeck, Germany), with a sensitivity of 90.3 and a specificity of 99.8% according to the manufacturer’s data. The maximum IgG level that could be determined with appropriate precision after dilution was 1920 relative units per milliliter (RU/ml).

#### SARS-CoV-2 Neutralization assay

Neutralizing antibodies were investigated using a live virus neutralization assay. A classical B.1.1.7 lineage (20I/501Y.V1) SARS-CoV-2 strain, previously isolated from a clinical specimen and propagated in Vero E6 cells, was used in all experiments. The whole genome sequence of the viral isolate was submitted to GISAID (accession reference EPI_ISL_1653931). In brief, serial 2-fold dilutions (starting from 1:10) of the heated serum (56°C for 30 min) were incubated for 1 h at 37°C with viral solution containing 100 TCID50 of SARS-CoV-2 and then added to Vero E6 cell monolayers in a 96-well plate. The cytopathic effect was recorded after 3 days, and the serum virus neutralization titer (V-NT50) was defined as the reciprocal value of the highest dilution that showed at least 50% protection of cells. A sample with a titer ≥ 20 was defined as positive. Negative signals were set to 0 for statistical analyses.

#### SARS-CoV-2 pseudovirus neutralization assay (pV-NT)

To further assess the neutralizing activity of sera, retroviral pseudoparticles containing the SARS-CoV-2 glycoprotein S (SARS-CoV-2pp) were produced as previously described (4), with a plasmid encoding the human "codon-optimized" sequence of the SARS-CoV-2 glycoprotein Spike (accession number: MN908947). The supernatants containing the SARS-CoV-2pp were harvested at 48-h post-transfection and filtered through a 0.45-μm membrane and stored at −80°C. The serum neutralization test was performed as previously described (5). In brief, 20 μl of SARS-CoV-2pp were incubated in the diluted serum at a final volume of 50 μl of DMEM+Glutamax+Penicillin-Streptomycin+10% Fetal Calf Serum (FCS) for 30 min at room temperature. The mixture was then added to HEK 293TT-ACE2 plated the day before (HEK 2932TT cells stably expressing the hACE2 receptor are seeded at 4500 cells/well in a volume of 50 μl of DMEM+Glutamax+Penicillin-Streptomycin+10% FCS mixture (6). At 48-h post-infection, Luciferase activity was measured using the Luciferase Assay System kit (Charbonnières-les-Bains, Promega FR) as recommended by the manufacturer and expressed as Relative Luciferase Units (RLUs). RLUs were compared and normalized to the wells where pseudoparticles were added in the absence of serum (100%). Serum pseudovirus neutralization titer 50 (pV-NT50) was expressed as the maximal dilution of the sera where the reduction of the signal is greater than 50%. The titer was multiplied by 781, since the initial volume of the sera tested was 8 μl and had to be normalized to 1 ml (7).

#### Peripheral blood mononuclear cells (PBMCs) preparation

Isolation and numeration of peripheral blood mononuclear cells (PBMCs) was performed from 10 to 15 ml of freshly collected heparinized blood samples. In brief, T cell Xtend (Oxford immunotec) at a concentration of 25 μl/ml of blood was added 15 min prior to isolation to remove cell debris and aggregates. SepMate-50 ml (StemCell Technologies) was then used for density gradient centrifugation. PBMCs were collected and washed twice using RPMI. Isolated cells were suspended in AIM-V medium and counted using flow cytometry with CD45 staining (Beckman Coulter) and Flow-Count fluorospheres (Beckman Coulter). Normalization of the cell suspension was performed at a final concentration of 2.5.106 cells/ml for T-CoV-Spot assay and 10.106 cells/ml for flow cytometry analyses.

#### IFNy ELISpot assay – T-CoV-Spot assay

T-CoV-Spot assay was performed as previously described (8). In brief, overlapping peptide pools covering the N-terminal S1 domain were used (PepTivator_ SARS-CoV-2, Miltenyi Biotec, Bergisch Gladbach, Germany). Peptides consisted of 15-mer sequences with 11 amino acids overlap. Microtitre plates coated with anti-IFNy antibodies (T-SPOT.TB, Oxford Immunotec) were used. The cell suspension was normalized at a final concentration of 2.5 × 10^6^ cells/ml, and plating with SARS-CoV-2 antigens was manually performed (2.5 × 105 PBMCs added per well). Peptide pools were added at a concentration of 0.5 μg/ml. Following an incubation at 37°C for 16–20 h in a humidified atmosphere containing 5% CO_2_, wells were washed and incubated with conjugate reagent for 1 h at 2–8°C. After a washing step, wells were developed for 7 min with substrate solution. The reaction was stopped by adding distilled water. Plates were allowed to dry in an oven at 37°C for 1 h. Spot-forming cells (SFCs) were detected using the CTL ImmunoSpot plate reader. Appropriate negative and positive controls were used (8).

#### Flow cytometry analyses

In addition to IFNγ secreting cells by ELISpot, SARS-CoV-2-specific T cell detection was also analyzed using flow cytometry. PBMC suspensions were normalized at a final concentration of 10×10^6^ cells/ml and 1×10.6 cells were incubated in RPMI for 16–20 h at 37°C in a humidified atmosphere containing 5% CO_2_. Then, 7-amino-actinomycin D (7AAD) (Biolegend), Pacific Blue-conjugated anti-CD107a antibody (clone H4A3; Beckman-Coulter) and the same peptides pools, at the same concentrations than for the ELISpot assay, were added to the cell suspension for 1 h (37°C, 5% CO_2_). Brefeldin A (Sigma-Aldrich) and monensin (Biolegend) were added at 2.5 microg/ml and 2 microM, respectively. The obtained cell preparation was conserved for 4 h (37°C, 5% CO_2_). The washed cells were then permeabilized with Cytofix/Cytoperm™ Fixation/Permeabilization Kit, according to the manufacturer recommendations (Becton Dickinson), and 2 washing steps with Perm/wash buffer were performed (Beckton Dickinson). For detection of surface molecules, antibodies against CD3 (APC-Alexa750 conjugated, clone UCHT1, Beckman Coulter), CD4 (APC, clone 13B8.2, Beckman Coulter), CD8 (Alexa700, clone B9.11, Beckman Coulter), CD154 (PE, clone TRAP-1, Beckman Coulter, IM2216U), CD69 (FITC, clone FN50, Biolegend, catalog no 310904) were used. Intracellular cytokines were detected with antibodies against TNFα (PC7, clone Mab11, Biolegend), IL2 (BV605, clone MQ1-17H12, Biolegend), IFNγ (BV650, clone 4S.B3, Biolegend). Each cell preparation was totally analyzed (around 300.000 T cells). For assessment of whole blood naive/memory T cells at baseline in LTCF residents, antibodies against CD4 (Pacific Blue, clone 13B8.2, Beckman Coulter), CD8 (APC, clone B9.11, Beckman Coulter, catalog no A99023), CD45RA (FITC, clone 2H4, Beckman Coulter) and CCR7 (PE, clone G043H7, Beckman Coulter) were used. Cells were analyzed on a Cytoflex S (Beckman Coulter) flow cytometer.

#### Fluorescence Activated Cell Sorter (FACS) data analysis

FACS data were analyzed with Kaluza Analysis Software (Beckman Coulter). The gating strategy for analysis of antigen-specific T cells is illustrated in the Extended Data Fig. 4). For the activation-induced markers (AIM) T cell assay, a specific T cell response was considered positive when the stimulation index was 2 or higher, *i.e*. when the antigen-stimulated cultures contained at least 2-fold higher frequencies of CD154+CD69+ cells among alive (7-AAD-) CD4+ T cells (AIM+ CD4+ T cells), or CD107a+CD69+ cells among alive CD8+ T cells (AIM+ CD8+ T cells), compared to the unstimulated control sample. No further background subtraction was applied. Co-expression of intracellular cytokines was assessed among AIM+ CD4+ and CD8+ T cells using a Boolean gating strategy. Unsupervised analysis was conducted using t-distributed stochastic neighbor embedding (t-SNE) in AIM+ CD4+ or CD8+ T cells (Cytobank, Beckman Coulter). All datasets were extracted from the pre-gating made with Kaluza on AIM+ T cells, group concatenations were made and all data were imported into Cytobank. Unsupervised cell subsets identification (clustering) was also performed for analysis of cytokines productions by AIM+ CD4+ and CD8+ T cells. Percentages of each main subsets of specific T cells (according to production of 0/1, 2 or 3 cytokines) obtained by the unsupervised FlowSOM analysis (considered as the addition of all clusters abundance in the subset) were reported on the subsets (Cytobank, Beckman Coulter).

#### Cytokine measurements

Plasma IL-1β, IL-6, TNFα and IL-10 concentrations were assessed using the Ella Automated Immunoassay System (ProteinSimple, San Jose, CA) following the manufacturer’s recommendations.

#### Statistical analyses

Categorical variables are expressed as numbers (percentages) and quantitative variables are expressed as median (interquartile range). Normality distribution was assessed graphically and using the Shapiro-Wilk test. Immune parameters were compared within the same group between the baseline and 3-month assessments using the Wilcoxon signed rank test. Comparisons of immune parameters between the four study groups (COVID-19-naive older, COVID-19-recovered older, COVID-19-naive young and COVID-19-recovered young) were done using the Kruskal–Wallis test followed by *post hoc* Dunn’s tests for quantitative measures and chi-squared test (or Fisher’s exact test in cases of expected cell frequency <5) for responder rates. Comparisons of baseline characteristics in COVID-19-recovered older adults and D90 characteristics in COVID-19-naive older adults (natural post-COVID-19 versus post-BNT162b2 immunization) were done using the Mann-Whitney U test. We assessed the correlation between age, vaccinal response parameters, nutritional, frailty or immunosenescence parameters by calculating Spearman’s rank correlation (r) coefficients, with their 95% confidence intervals based on the Fisher Z-transformation. Statistical tests were done at the two-tailed α level of 0.05. No correction for multiple testing was carried out. Data analyses and graphs were performed using the GraphPad Prism software version 9.1.2 (GraphPad Software, La Jolla, CA, USA).

#### Ethics

This study was performed in accordance with the Declaration of Helsinki principles for ethical research. The study was approved by the Ile-De-France V (ID-CRB 2021-A00119-32) ethics committee. All participants (and/or their legal representative if required) received detailed information and signed a consent form before participating in the study.

The study was registered in ClinicalTrials.gov, with the identifier NCT04760704.

---

**Supplementary Table 1.**
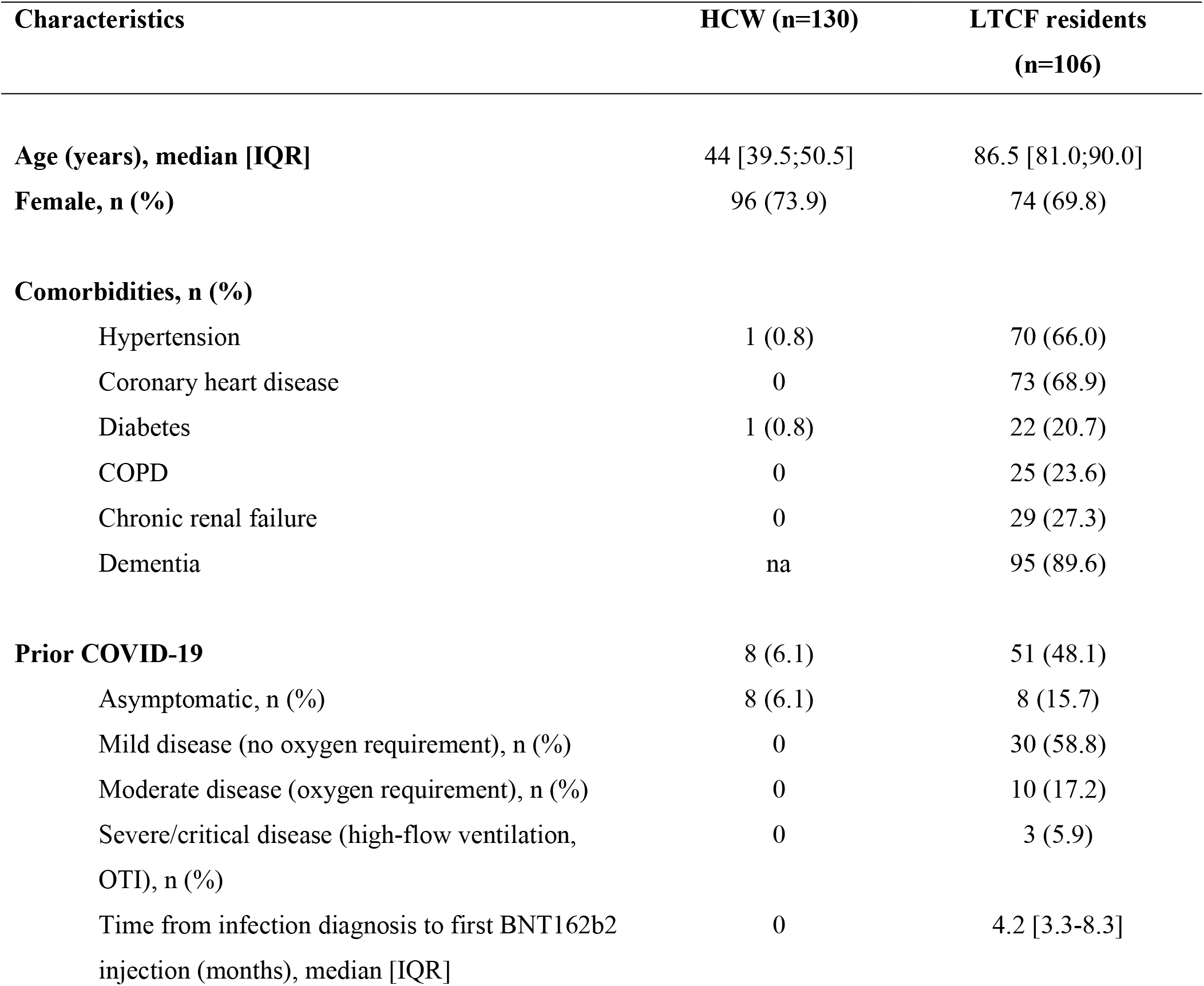
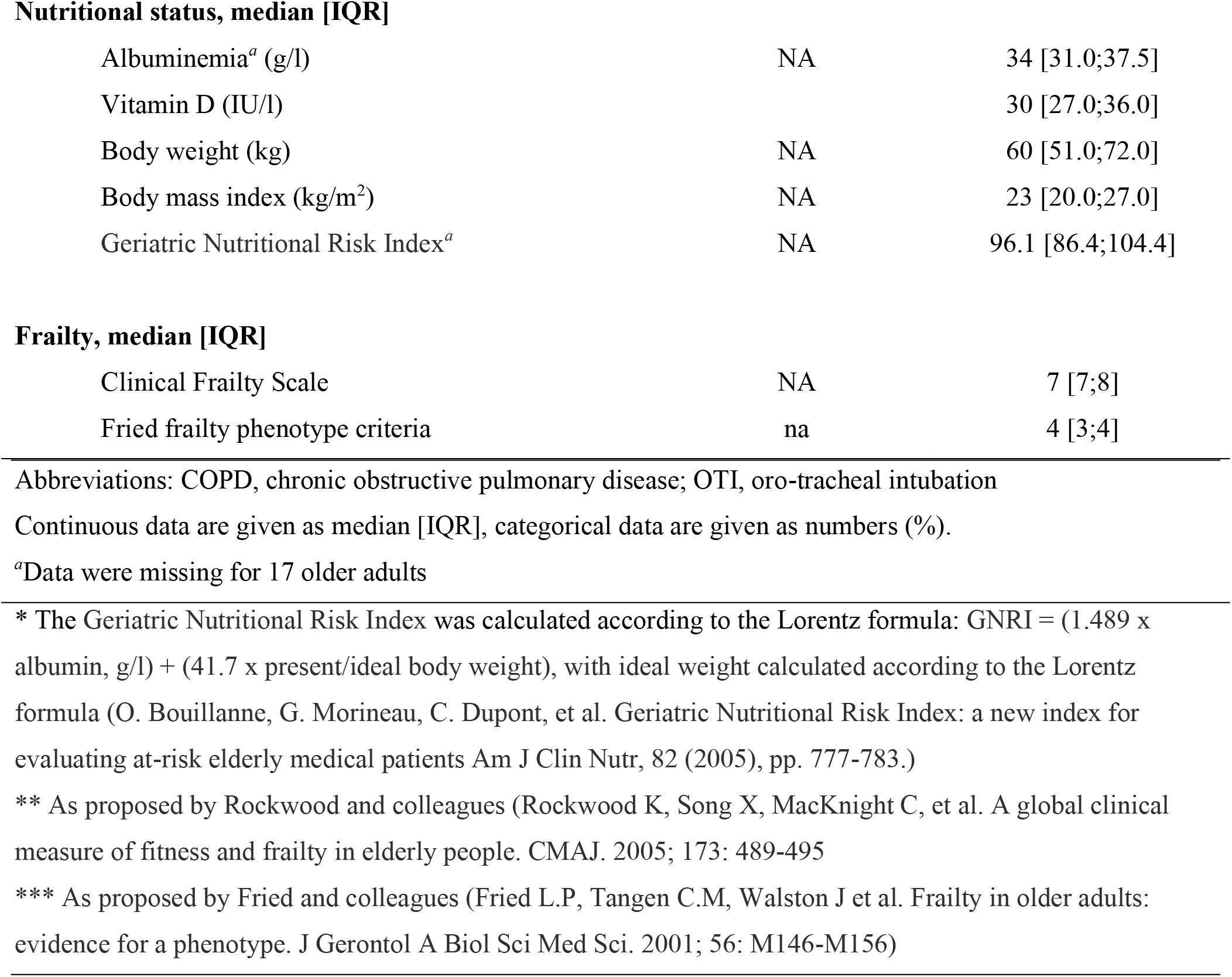
Characteristics of young and older adults.

| Characteristics | HCW (n=130) | LTCF residents<br>(n=106) |
| --- | --- | --- |
| **Age (years), median [IQR]** | 44 [39.5;50.5] | 86.5 [81.0;90.0] |
| **Female, n (%)** | 96 (73.9) | 74 (69.8) |
| **Comorbidities, n (%)** |  |  |
| Hypertension | 1 (0.8) | 70 (66.0) |
| Coronary heart disease | 0 | 73 (68.9) |
| Diabetes | 1 (0.8) | 22 (20.7) |
| COPD | 0 | 25 (23.6) |
| Chronic renal failure | 0 | 29 (27.3) |
| Dementia | na | 95 (89.6) |
| **Prior COVID-19** | 8 (6.1) | 51 (48.1) |
| Asymptomatic, n (%) | 8 (6.1) | 8 (15.7) |
| Mild disease (no oxygen requirement), n (%) | 0 | 30 (58.8) |
| Moderate disease (oxygen requirement), n (%) | 0 | 10 (17.2) |
| Severe/critical disease (high-flow ventilation, OTI), n (%) | 0 | 3 (5.9) |
| Time from infection diagnosis to first BNT162b2 injection (months), median [IQR] | 0 | 4.2 [3.3-8.3] |
| **Nutritional status, median [IQR]** |  |  |
| Albuminemia<sup>a</sup> (g/l) | NA | 34 [31.0;37.5] |
| Vitamin D (IU/l) |  | 30 [27.0;36.0] |
| Body weight (kg) | NA | 60 [51.0;72.0] |
| Body mass index (kg/m<sup>2</sup>) | NA | 23 [20.0;27.0] |
| Geriatric Nutritional Risk Index<sup>a</sup> | NA | 96.1 [86.4;104.4] |
| **Frailty, median [IQR]** |  |  |
| Clinical Frailty Scale | NA | 7 [7;8] |
| Fried frailty phenotype criteria | na | 4 [3;4] |

Abbreviations: COPD, chronic obstructive pulmonary disease; OTI, oro-tracheal intubation
Continuous data are given as median [IQR], categorical data are given as numbers (%).
<sup>a</sup>Data were missing for 17 older adults
\* The Geriatric Nutritional Risk Index was calculated according to the Lorentz formula: GNRI = (1.489 x albumin, g/l) + (41.7 x present/ideal body weight), with ideal weight calculated according to the Lorentz formula (O. Bouillanne, G. Morineau, C. Dupont, et al. Geriatric Nutritional Risk Index: a new index for evaluating at-risk elderly medical patients Am J Clin Nutr, 82 (2005), pp. 777-783.)
\*\* As proposed by Rockwood and colleagues (Rockwood K, Song X, MacKnight C, et al. A global clinical measure of fitness and frailty in elderly people. CMAJ. 2005; 173: 489-495
\*\*\* As proposed by Fried and colleagues (Fried L.P, Tangen C.M, Walston J et al. Frailty in older adults: evidence for a phenotype. J Gerontol A Biol Sci Med Sci. 2001; 56: M146-M156)

**Supplementary Table 2.** Significant correlations between age and main immune parameters of the post vaccinal response at 3 months.

| Correlation | Spearman's rank (r) | P value | 95% CI | Sample size |
| --- | --- | --- | --- | --- |
| Age with |  |  |  |  |
| Anti-S1 IgG levels | -0.36 | < 0.0001 | -0.48;-0.22 | 181 |
| NT50 LV-NT | -0.35 | < 0.0001 | -0.48;-0.19 | 146 |
| NT50 pV-NT | -0.56 | < 0.0001 | -0.66;-0.44 | 160 |
| S1 reactive T cells | -0.25 | 0.0008 | -0.38;-0.10 | 179 |
| (ELISpot) |  |  |  |  |
| AIM+ CD8+ (index) | -0.23 | 0.004 | -0.37;-0.07 | 159 |
| Anti-S1 IgG antibodies with |  |  |  |  |
| NT50 LV-NT | 0.79 | < 0.0001 | 0.72;0.85 | 181 |
| NT50 pV-NT | 0.68 | < 0.0001 | 0.58;0.75 | 146 |
| S1 reactive T cells | 0.45 | < 0.0001 | 0.32;0.56 | 160 |
| (ELISpot) |  |  |  |  |
| AIM+ CD4+ (index) | 0.21 | 0.007 | 0.05;0.36 | 179 |
| AIM+ CD8+ (index) | 0.20 | 0.001 | 0.05;0.35 | 159 |
| NT50 LV-NT with |  |  |  |  |
| NT50 pV-NT | 0.65 | < 0.0001 | 0.53;0.74 | 127 |
| S1 reactive T cells | 0.44 | < 0.0001 | 0.29;0.56 | 145 |
| (ELISpot) |  |  |  |  |
| NT50 pV-NT with |  |  |  |  |
| S1 reactive T cells | 0.39 | < 0.0001 | 0.25;0.52 | 158 |
| (ELISpot) |  |  |  |  |
| AIM+ CD4+ (index) | 0.29 | 0.0004 | 0.13;0.44 | 140 |
| AIM+ CD8+ (index) | 0.25 | 0.002 | 0.09;0.4 | 140 |
| AIM+ CD4+ (index) with |  |  |  |  |
| AIM+ CD8+ (index) | 0.21 | 0.009 | 0.05;0.35 | 159 |

## SUPPLEMENTARY FIGURES

### Extended Data

**Extended Data Fig. 1:**
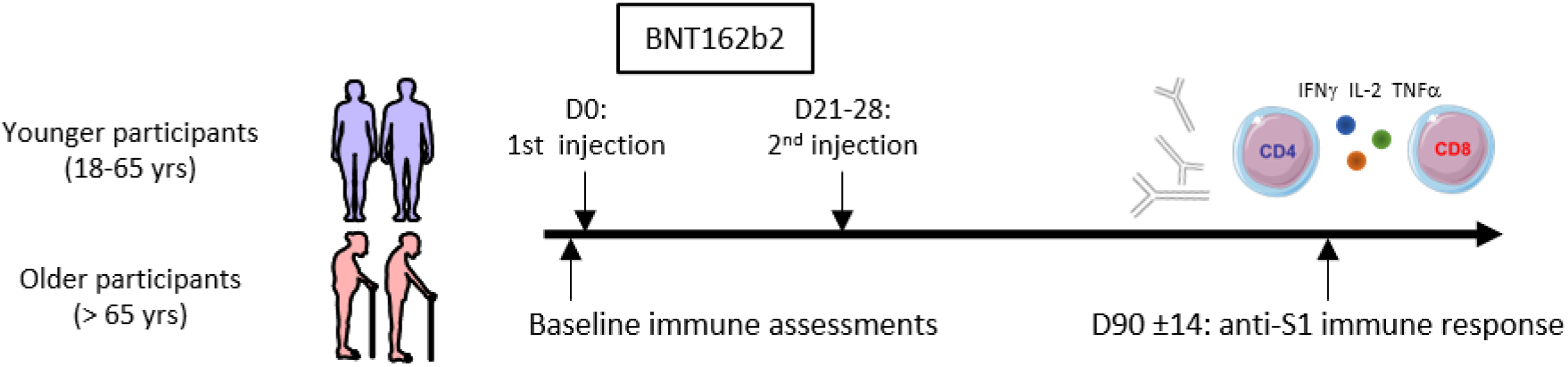
General study design.

**Extended Data Fig. 2:**
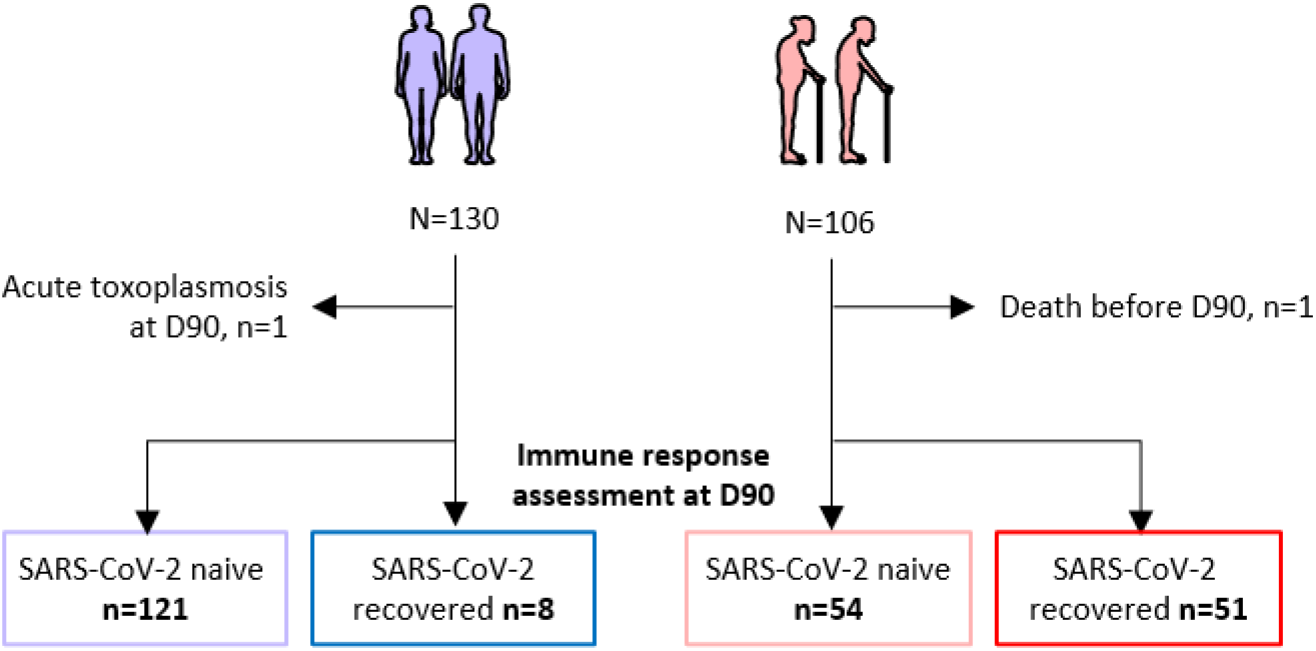
Flow chart of the study.

**Extended Data Fig. 3:**
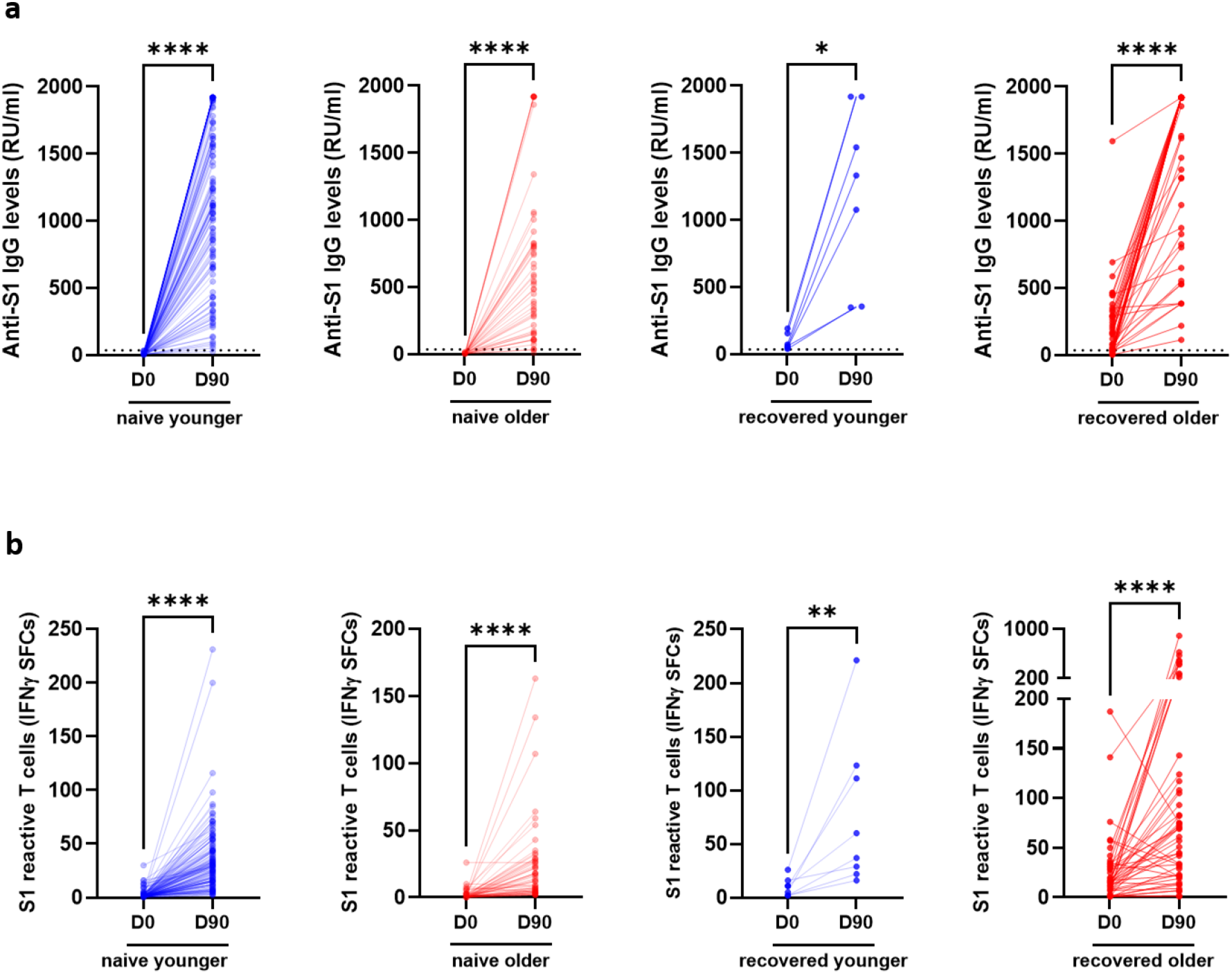
Anti-S1 IgG and S1 reactive T cells (ELISpot) at D0 and D90. Wilcoxon matched-pairs signed rank test was used for paired comparisons. *P* values * <0.05, ** <0.01, **** <0.0001. CTL, IFNγ SFCs, interferon gamma spot forming cells.

**Extended Data Fig. 4:**
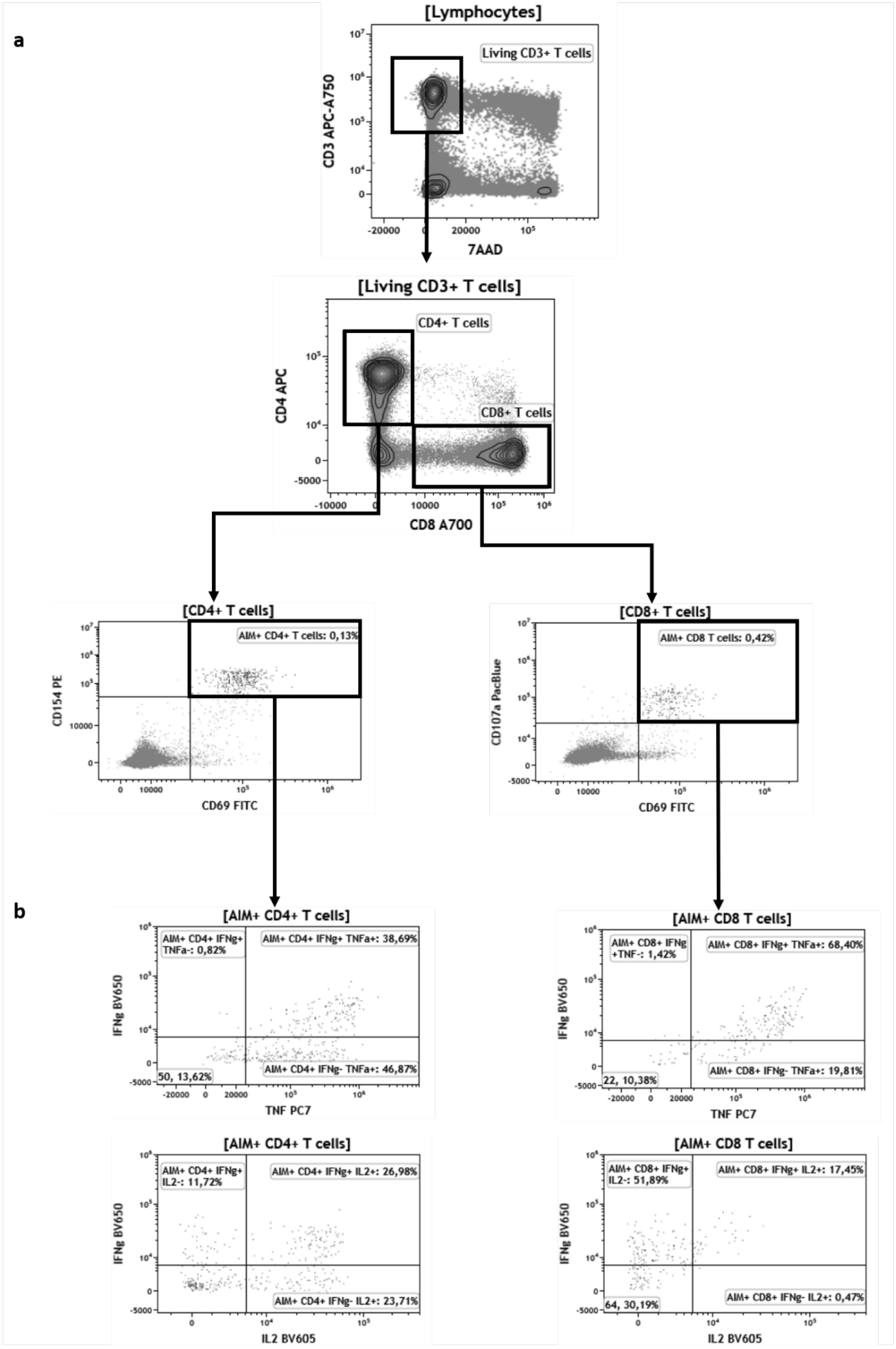
Gating strategy for flow cytometry analyses of CD4+ and CD8+ T cells after BNT162b vaccination. **a**, Identification of activation induced markers (AIM+ cells). Briefly, “living CD3+ T cells” are identified as 7AAD (7-aminoactinomycine D) negative and CD3 positive cells. Among this population, CD4+ and CD8+ T cells are selected according to CD4+ and CD8+ expression, respectively. AIM+ cells among CD4+ T cells are both CD154+ and CD69+. AIM+ cells among CD8+ T cells are both CD107a+ and CD69+. **b**, Representative plots displaying IFNγ, IL-2 and TNFα expression among AIM+ CD4+ and CD8+ T cells.

**Extended Data Fig. 5:**
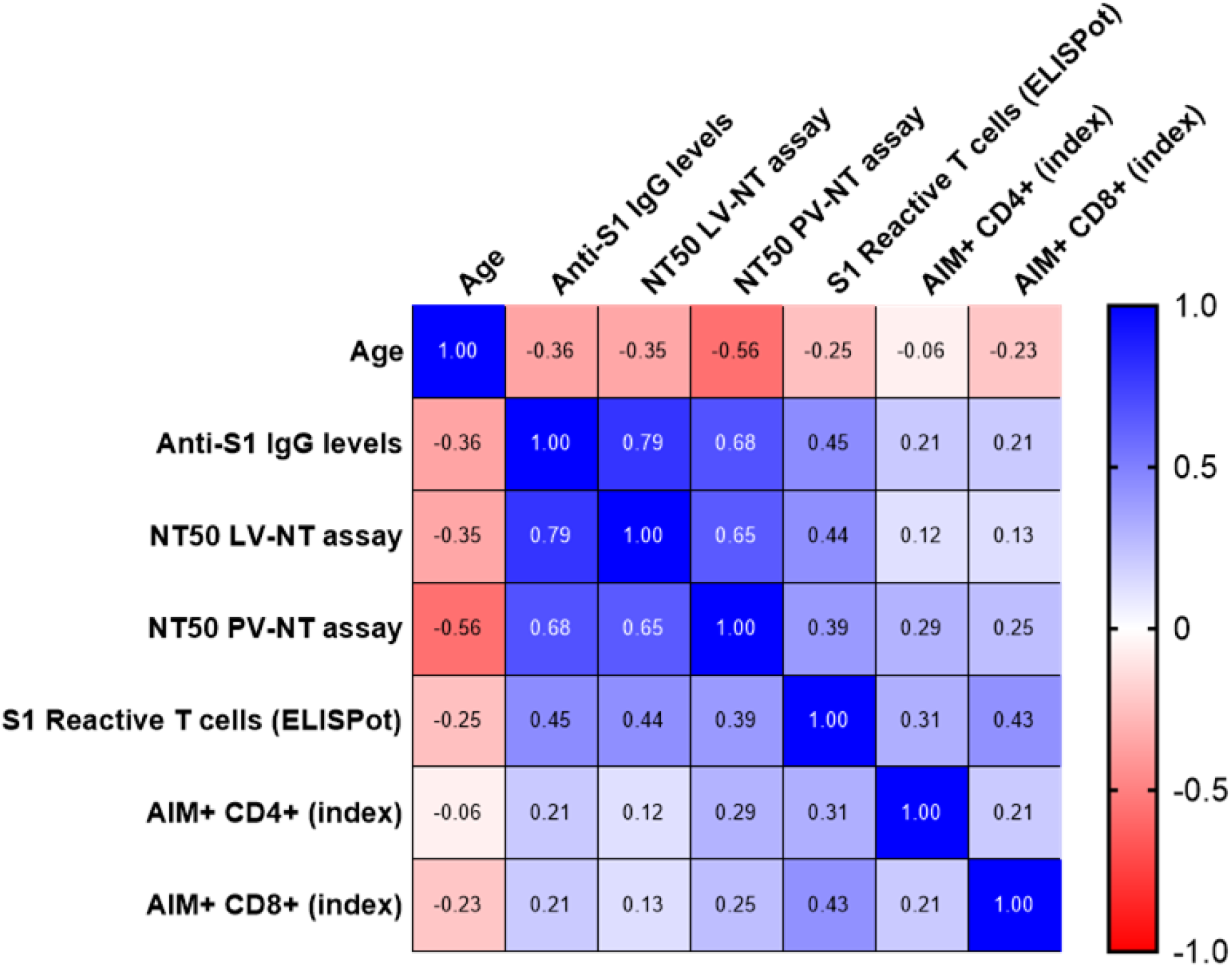
Correlations between age and main immune parameters of the post vaccinal response at 3 months. Values are Spearman’s rank correlation (r) coefficients. The number of pairs that were analyzed, *P* values and 95% confidence intervals of significant correlations are detailed in Supplementary Table 2. AIM^+^, cell expressing activation induced markers; NT50 LV-NT assay, 50% serum neutralization titer in live virus neutralization assay; NT50 PV-NT assay, 50% serum neutralization titer in pseudovirus neutralization assay.

**Extended Data Fig. 6:**
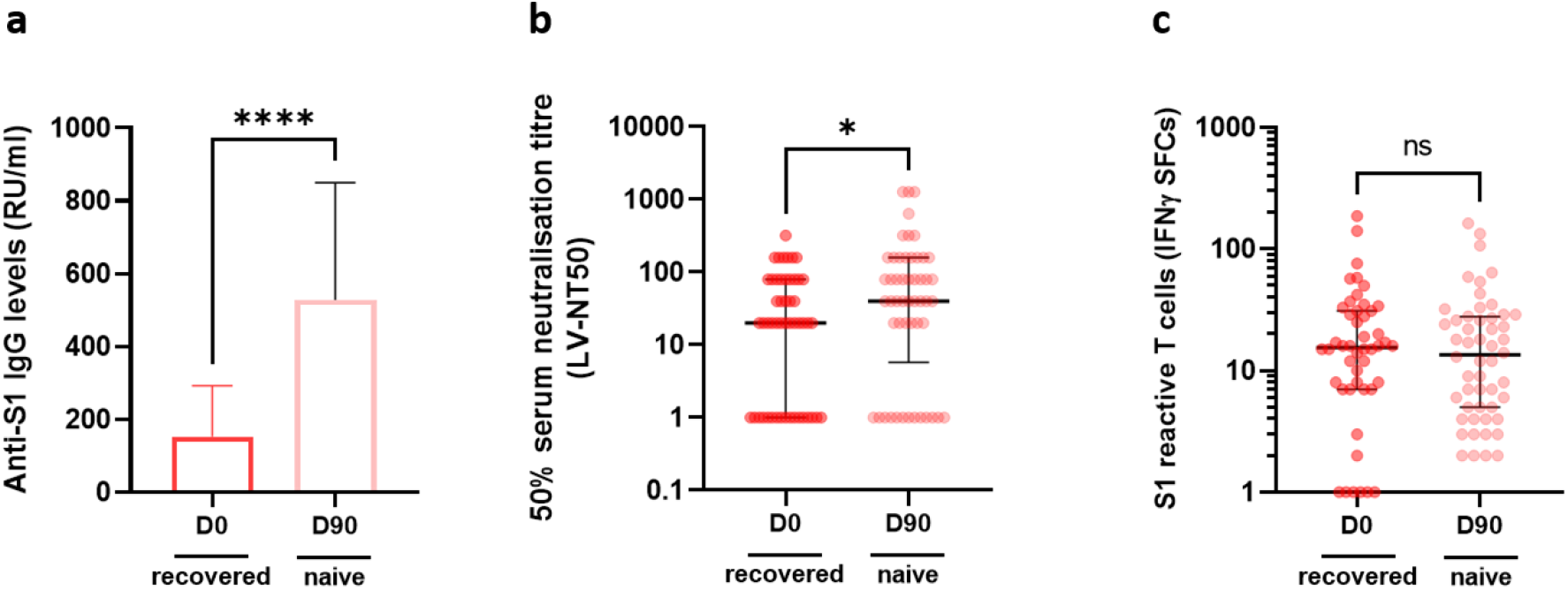
Specific antibody and T cell responses after COVID-19 and after BNT162b2 in elderly LTCF residents. **a,** antibody responses assessed by ELISA (anti-S1 IgG) (COVID-19-naive older *n* = 54, COVID-19-recovered older *n* = 49). **b**, serum neutralization assay against live virus (COVID-19-naive older *n* = 51, COVID-19-recovered older *n* = 52). **c**, number of S1 peptide pool reactive T cells (ELISpot) (COVID-19-naive older *n* = 52, COVID-19-recovered older *n* = 48). Conval D0, value at baseline, prior BNT162b2, in COVID-19-recovered older participants, IFNγ SFCs, interferon gamma spot forming cells; LV-NT50, 50% serum neutralization titer in live virus neutralization assay; Naive D90, value at 3 months after first injection of BNT162b2, in COVID-19-naive older participants.

**Extended Data Fig. 7:**
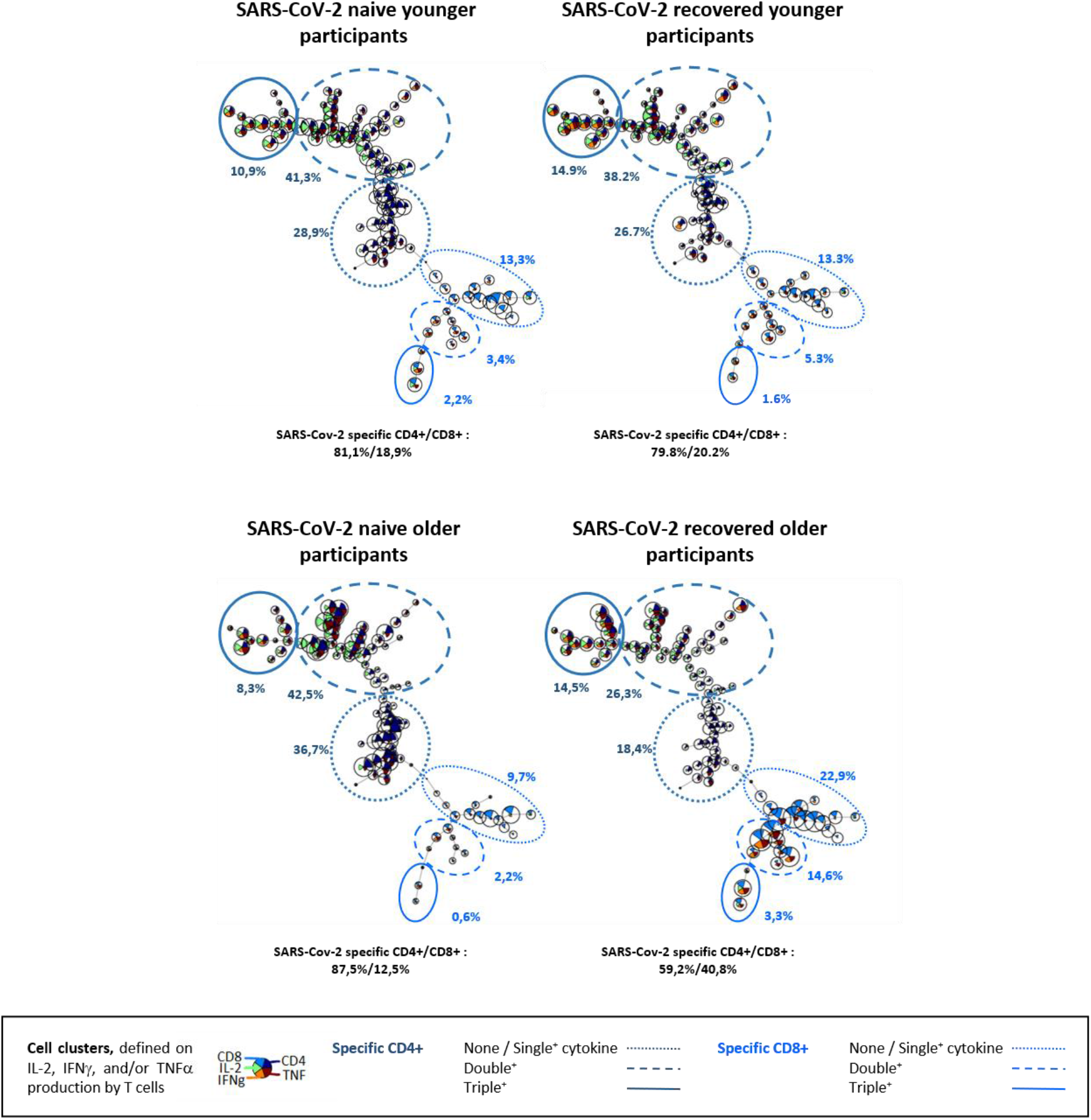
Cluster analysis of specific T cells subset after BNT162b2 in elderly LTCF residents and in HCWs (controls). FlowSOM results for COVID-19-naive and COVID-19-recovered young adults (top) and for COVID-19-naive and COVID-19-recovered older adults (bottom). Cell clusters were defined according to IL-2, IFNγ and TNFα expression. Manual metaclusters were identified among specific CD4+ T cells (dark blue) and specific CD8+ T cells (light blue) for cells producing none or one (small dotted line), two (large dotted line) or three cytokines (plain line) out of INFγ, IL-2 and TNFα.

**Extended Data Fig. 8:**
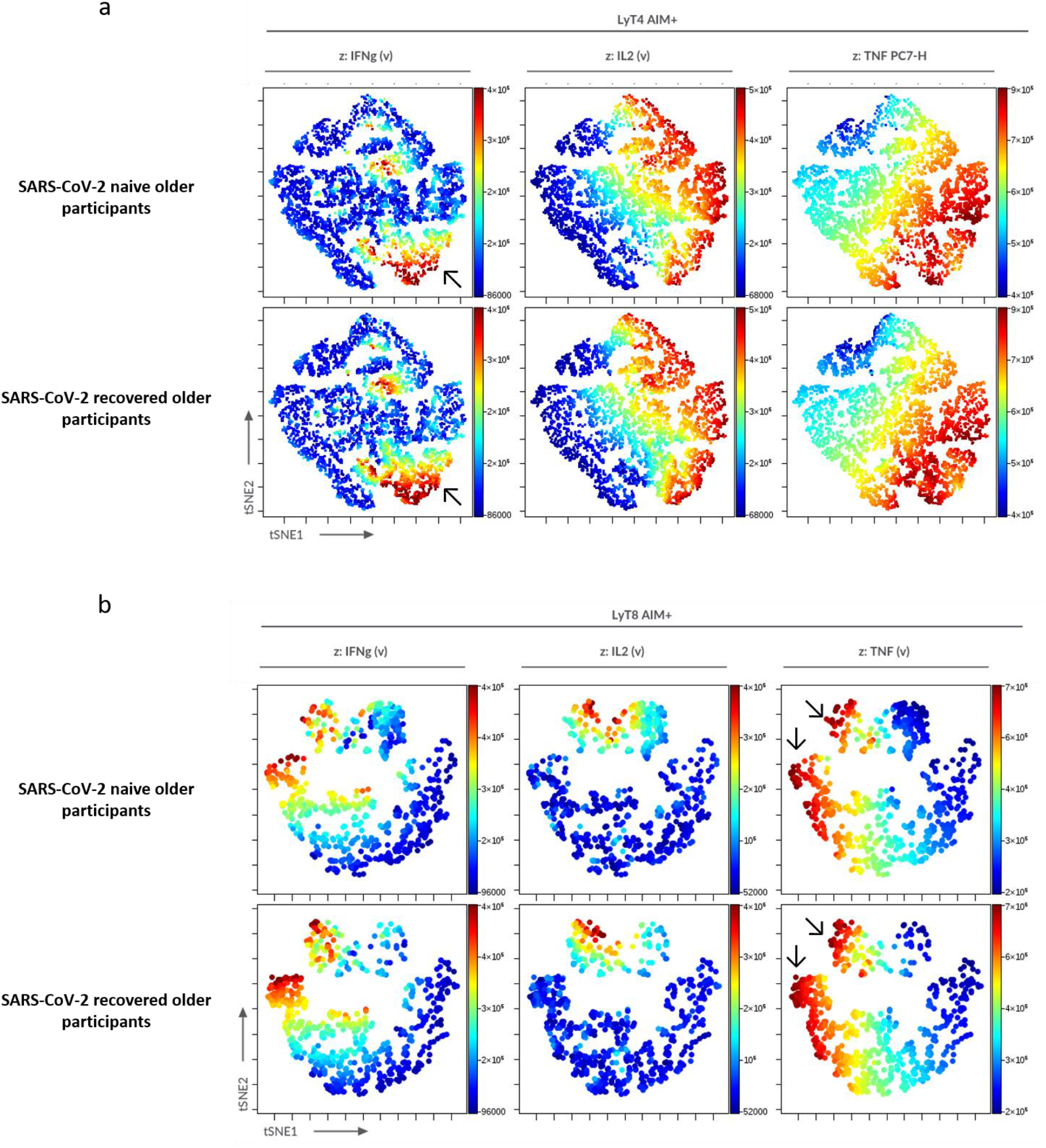
Unsupervised analysis of CD4+ and CD8+ T cell functionality in elderly LTCF residents using tSNE. **a.** AIM+ CD4+ T cells from all groups were concatenated and subjected to unsupervised analysis using t-distributed stochastic neighbor embedding (t-SNE); highlighted (z-dimension) are areas with IFNγ, IL-2 or TNFα cell expression in COVID-19-naive and COVID-19-recovered older adults. To be noted, the higher frequency of IFNγ+ CD4+ T cells in COVID-19-naïve older adults (arrow). **b.** AIM+ CD8+ T cells from all groups were concatenated and subjected to unsupervised analysis using t-distributed stochastic neighbor embedding (t-SNE); highlighted (z-dimension) are areas with IFNγ, IL-2 or TNFα cell expression in COVID-19-naive and COVID-19-recovered older adults. To be noted, the higher frequency of TNFα+ CD8+ T cells in COVID-19-recovered older adults (arrow).

**Extended Data Fig. 9:**
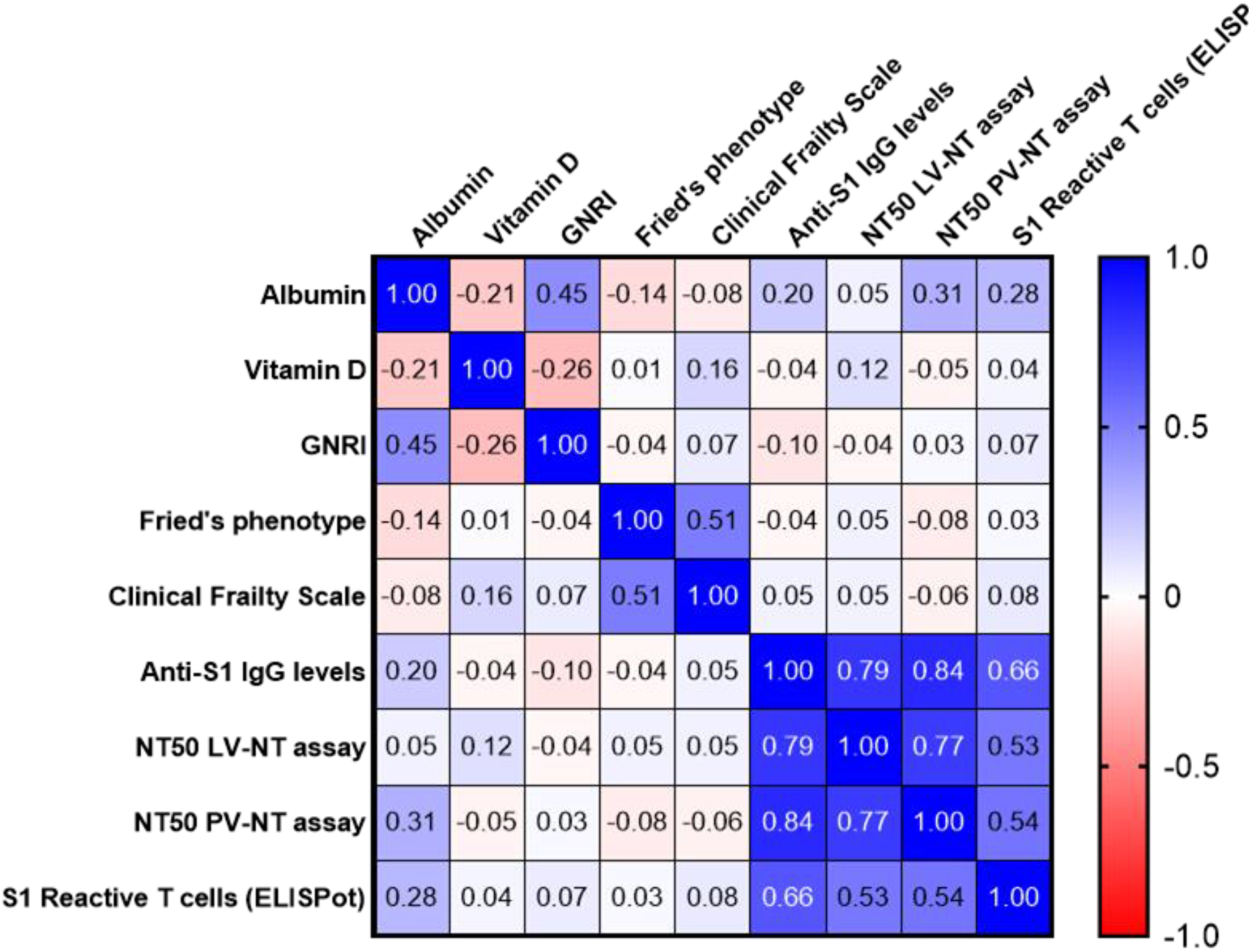
Correlations between nutritional status and frailty scale at baseline, and main immune parameters of the post vaccinal response at 3 months. The values correspond to Spearman’s rank correlation (r) coefficients. Only one correlation was found significant, between Albumin level and NT50 pV-NT (*r [95% CI)* 0.31 [0.022;0.55], *P* = 0.031, sample size *n* = 49). AIM^+^, cell expressing activation induced markers; NT50 LV-NT assay, 50% serum neutralization titer in live virus neutralization assay; NT50 pV-NT assay, 50% serum neutralization titer in pseudovirus neutralization assay.

**Extended Data Fig. 10:**
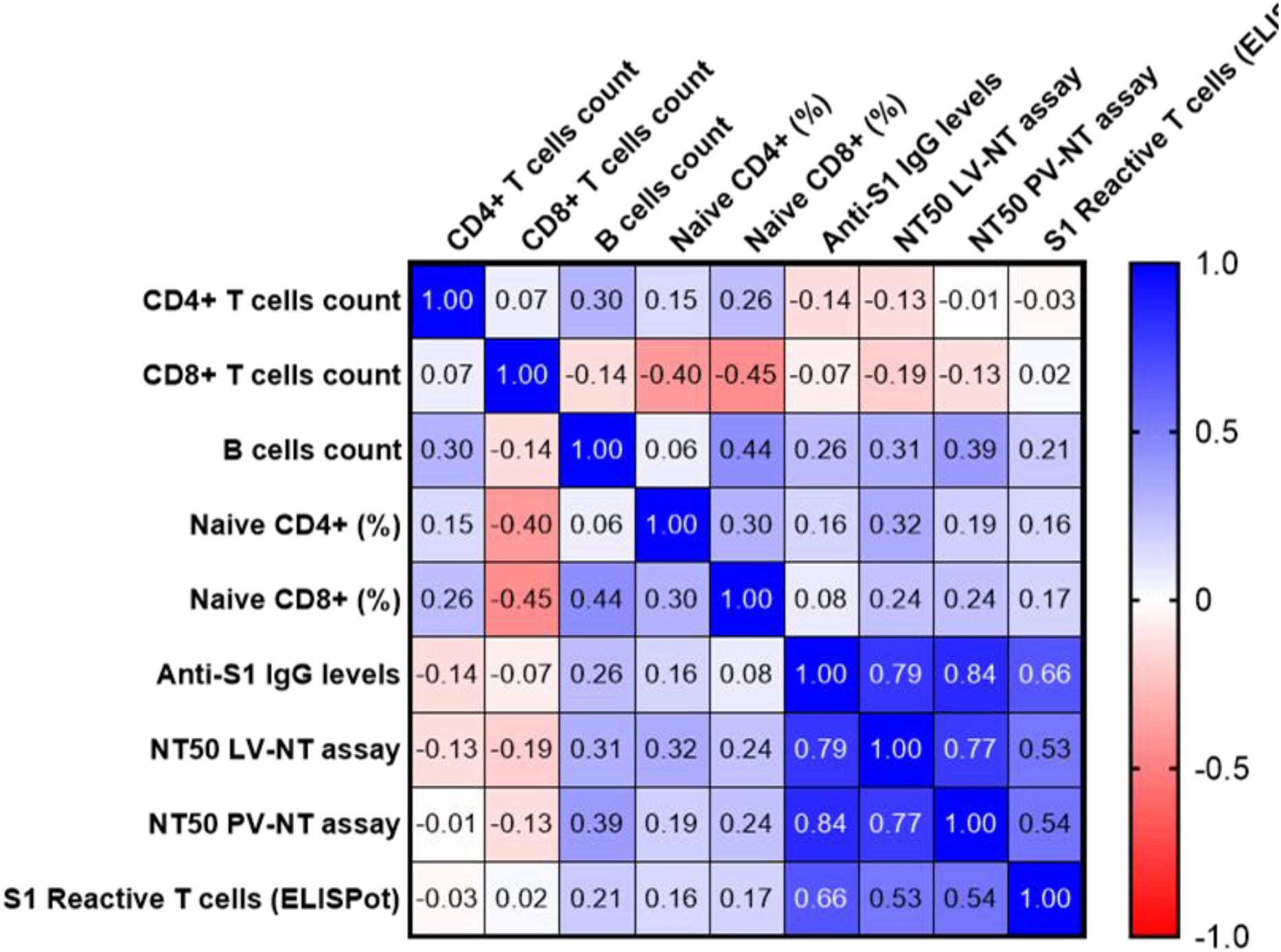
Correlations between T and B cells counts at baseline, and main immune parameters of the post vaccinal response at 3 months. The values correspond to Spearman’s rank correlation (r) coefficients. Only one correlation was found to be significant, between B cell count level and NT50 pV-NT (*r [95%CI)* 0.39 [0.009;0.63], *P* = 0.012, sample size *n* = 41). AIM^+^, cell expressing activation induced markers; NT50 LV-NT assay, 50% serum neutralization titer in live virus neutralization assay; NT50 PV-NT assay, 50% serum neutralization titer in pseudovirus neutralization assay.

**Extended Data Fig. 11:**
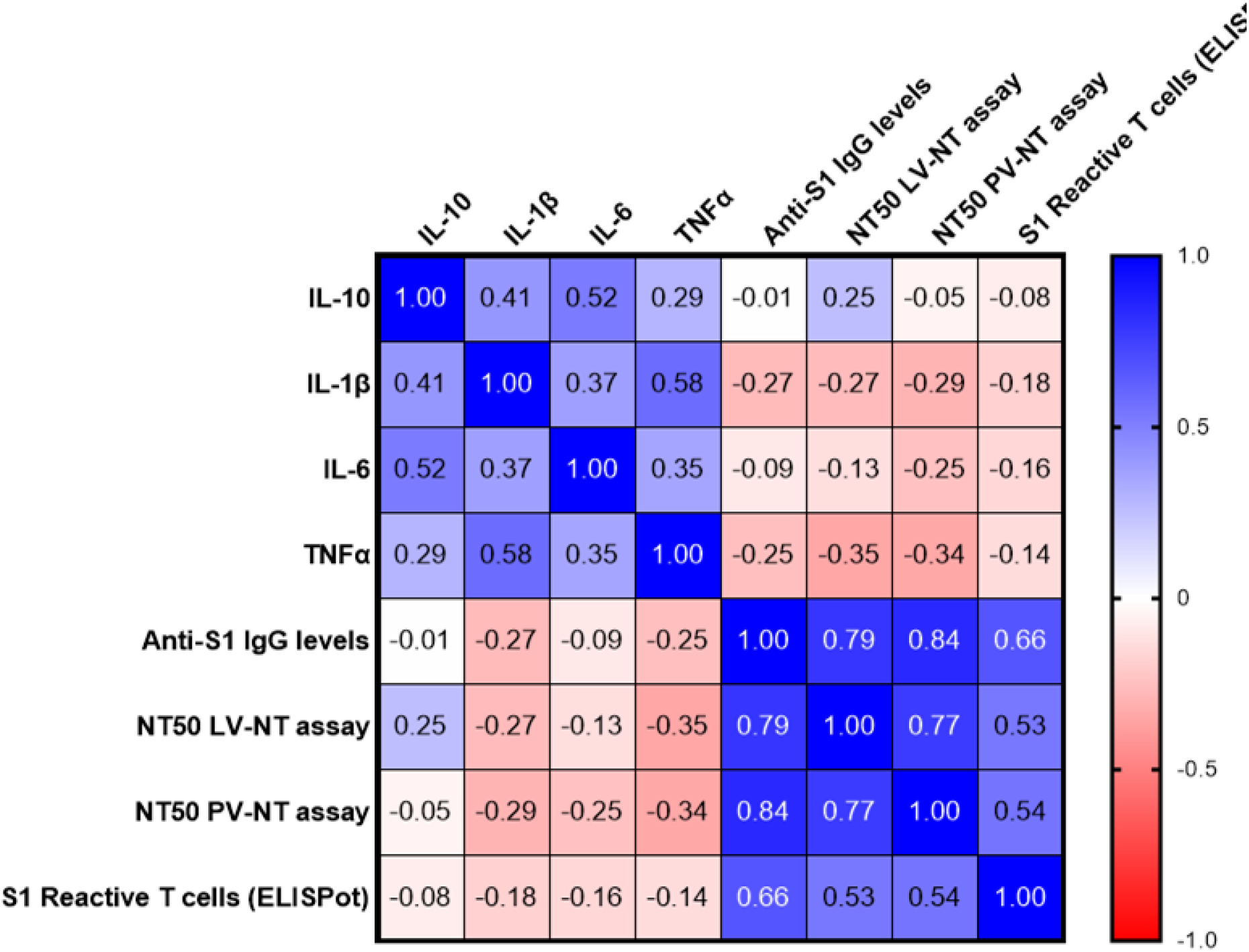
Correlations between plasma cytokines levels at baseline, and main immune parameters of the post vaccinal response at 3 months. The values correspond to Spearman’s rank correlation (r) coefficients. Only 2 correlations were found to be significant, between TNFα levels and NT50 LV-NT (*r [95% CI)* −0.35 [−0.62;0.007], *P* = 0.048, sample size *n* = 33), and between TNFα levels and NT50 pV-NT (*r [95%CI)* −0.34 [-0.60;0.02], *P* = 0.034, sample size *n* = 38). AIM^+^, cell expressing activation induced markers; NT50 LV-NT assay, 50% serum neutralization titer in live virus neutralization assay; NT50 PV-NT assay, 50% serum neutralization titer in pseudovirus neutralization assay.

## Notes

### Competing Interest Statement

The authors have declared no competing interest.

